# Discrepancies in Biomarker Identification in Different Peak-Picking Strategies in Untargeted Metabolomics Analyses of Cells, Tissues, and Biofluids

**DOI:** 10.1101/2025.03.04.641559

**Authors:** Iqbal Mahmud, Taylor A. Harmon, Laurel E. Meke, Timothy J. Garrett

**Affiliations:** Department of Pathology, Immunology, and Laboratory Medicine, University of Florida College of Medicine, Gainesville, FL 32610; Department of Chemistry, College of Liberal Arts and Sciences, University of Florida, Gainesville, FL 32611; Southeast Center for Integrated Metabolomics (SECIM), Clinical and Translation Research Institute, University of Florida, Gainesville, FL 32610; Department of Bioinformatics and Computational Biology, The University of Texas MD Anderson Cancer Center, Houston TX, USA

**Keywords:** Untargeted Metabolomics, Peak picking, XCMS, MZmine, Biomarker

## Abstract

Different software and algorithms are available for peak picking in nontargeted metabolomics and each may have its own strengths and limitations. The choice of pick picking method can significantly influence the results obtained, including the number and identity of metabolites detected, their quantification, and subsequent biomarker analysis. The impact of peak picking by different tools in an untargeted metabolomics-based biomarker study is largely understated. This study compares two popular open-source software tools for peak picking in untargeted metabolomics of cancer cells, tissues, and bio-fluids: XCMS and MZmine 2. The investigation evaluates the impact of these peak-picking algorithms on biomarker identification. We found significant discrepancy between the results obtained from XCMS and MZmine 2, regardless of the sample types, solvent gradient phases, retention time (RT), or mass-to-charge ratio (m/z) tolerances used. Notably, this study revealed significant disagreement between peak picking tools in the context of metabolite-based biomarker study and highlights the importance of carefully evaluating and selecting appropriate peak picking tools to ensure reliable and accurate results in untargeted metabolomics research.

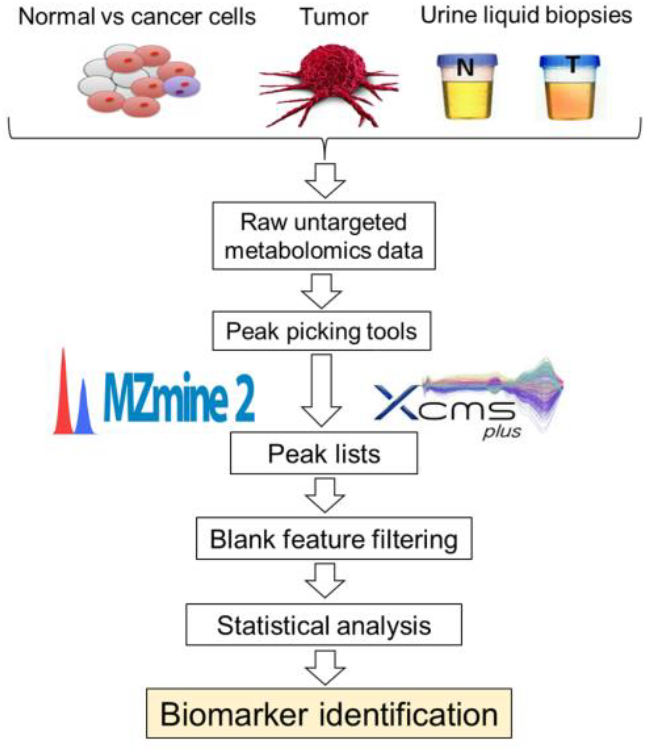

## INTRODUCTION

Untargeted metabolomics has become a popular technique with the aim of characterizing biochemical differences between phenotypically distinct populations.^1–6^ Significant developments in analytical instrumentation, particularly the advancements in both mass and chromatographic resolution over the past two decades, have continued to improve the quality and the information density of metabolomic data.^7,8^ Because of these improvements, the increasing complexity of experimental data sets has necessitated the use of peak-picking algorithms.^9^

In most software, extraneous or false peaks can severely limit the utility of collected data for biomarker identification.^10,11^ Several efforts, including blank feature filtering (BFF)^12^, peak annotation and verification engine (PAVE)^13^, and system-level annotation (creDBle)^7^, have improved the efficiency of metabolic peak picking in untargeted metabolomics. However, the bias of peakpicking packages has not been thoroughly investigated in regard to biomarker identification.

XCMS and MZmine 2 are two of the most commonly used data preprocessing tools for untargeted metabolomics analysis.^9,14^ One group previously demonstrated differences in overall discrimination and selectivity between MZmine 2 and XCMS.^9^ Several other studies have also compared metabolic peak picking among several freeware and commercial software packages.^14,15^ For instances, similarity among several peak picking software was studied using a 10 ppm *m/z* window and 0.15 min difference in retention time^15^. Using a mixture of 84 standard compounds, the researchers identified 24 metabolites in positive mode and 13 in negative mode. Only 13 and 8 standard metabolites were detected as overlapping among the four software packages (PeakView®, MarkerView**™**, MetabolitePilot**™**, and XCMS online) in positive and negative mode, respectively^15^. Another study found similar discrepancies between MZMine 2 and XCMS in analyses of human plasma^9^. Most recently, different untargeted metabolome processing tools such as MZmine2, enviMass, Compound Discoverer, and XCMS online results also showed low coherence among intersecting metabolic features in environmental samples.^16^ Interestingly, Myers et al. identified several problems in the peak detection algorithm of MZMine and XCMS.^9^ Failure to properly separate, detect, and identify suitable metabolic biomarkers is a major issue in bioanalytical chemistry^17,18^ and hence, it demands in-depth elucidation of metabolic features.

Peak detection in untargeted metabolomics plays a crucial role in the elucidation of biomarkers. It directly affects the quality and reliability of metabolomics data. Therefore, evaluating the reliability, reproducibility, and transferability of results obtained from different software packages is of paramount importance. Herein, we analyzed three different biological sample sets using MZmine 2 and XCMS.

Different statistical tools such as principal component analyses (PCA) and partial least squares-discriminate analyses (PLS-DA) were utilized to evaluate the bioanalytical impact of peak-picking discrepancies. Receiver operating curve (ROC) and Venn diagrams were employed to diagnose the specific causes of observed differences between MZmine 2 and XCMS. Biomarkers were annotated and mapped with METLIN and KEGG databases, respectively. Although biomarker discovery and identification continue to be a major problem in metabolomics workflows, this investigation seeks to uncover disparities between the two most widely used metabolic peak-picking software packages: MZmine2 and XCMS. By identifying these disparities, it may be possible to improve the accuracy and reliability of metabolomics analyses. In recent years, non-targeted metabolomics and lipidomics have gained attention, and they have been found to complement upstream transcriptomics, proteomics, and genomics data.^19–24^ This integration of different omics datasets can provide a more comprehensive understanding of biological mechanism and enable the identification of meaningful biomarker. Furthermore, stable isotope tracers can be used to track the flow of labeled metabolites through various biochemical reactions or metabolic pathways.^25–27^ By incorporating stable isotope tracers into global metabolomics experiment, the data quality can be improved^28^, as it provides insights into the dynamic behavior of metabolites and their transformations within the cellular system.

## EXPERIMENTAL SECTION

### Thyroid Tissue

Thyroid tissue was supplied by the Biorepository at the Clinical and Translational Sciences Institute at the University of Florida. Sample comprised ∼0.5 cm^3^ sections each of resected nodule and normal gland from three patients.

### Cell

Normal oral cell, periodontal ligament fibroblast (PLDF) and oral tumor cell, OQ01 were supplied by Dr. Ann Progulske-Fox lab in the department of Oral Biology at the University of Florida. Each group comprised triplicate cell pellets where each replicate contained one million cells.

### Urine

Urine samples from 10 prostate cancer (PCa) patients were obtained from the Biospecimen Pathology Core of the SPORE in Prostate Cancer at Northwestern University, Robert H. Lurie Comprehensive Cancer Center (P50 CA180995). Urine samples from 10 healthy individuals with no prior medical history of cancer were obtained from the Life Study (U01AG022376, University of Florida, Gainesville FL). The institutional review board (IRB) approved the sample use. Urine samples were collected using urine preservation tubes (Norgen Bioteck, Thorold, ON) and stored at −80 °C until further use for UHPLC−HRMS analyses.

### Global metabolite extraction for tissue

All samples were given as 1 cm^3^ cubes of resected tissue. The entire sample was homogenized by bead beater four times in 50 μL of 5 mM ammonium acetate. Each homogenized sample was centrifuged at 20,000 rcf for 20 min at 4 °C and collected 35 μL of supernatant from each homogenate. Each sample was analyzed by Qubit fluorimetry and diluted to 500 μg/mL total protein concentration. To 50 μL of this diluted solution was added 400 μL 8:1:1 acetonitrile:methanol:acetone. Samples were votexed, 10 s, and incubated for 10 min at 4 °C to further precipitate proteins, then centrifuged at 20,000 rcf for 20 min at 4 °C. The supernatant (375 μL) was collected and dried down under low heat (30 °C) and nitrogen.

### Global metabolite extraction for cells

Global metabolomics of cells were conducted for 1 million cells per replicate. Briefly, cell samples were homogenized using a bead beater. Cells were prenormalized to a protein concentration of 300 μg/mL as measured by Qubit® fluorescence assay. Protein precipitation was performed using 1 mL ice-cold methanol (80%) for 30 minutes with occasional vortexing. The samples were centrifuged at 20,000 rpm for 10 minutes at 4 °C to pellet the protein. The supernatant was transferred to a new tube and dried under nitrogen gas at 30 °C.

### Global metabolite extraction for Urine

In case of urine, 100 μL of raw urine was utilized for subsequent extraction and precipitation as previously discussed.^6,29^ Protein precipitation was performed using 1 mL ice-cold methanol (80%) for 30 minutes with occasional vortexing. Sample solution was centrifuged as described above, and 500 μL of supernatant was transferred to fresh tubes. Samples were dried under nitrogen and 30 °C heat before reconstitution in 50 μL 0.1 % formic acid.

### Metabolite extraction monitor

Throughout the metabolite extraction of tissue, cells, and urine samples, 20 μL of a metabolite internal standard mixture containing L-leucine-D10 (4μg/mL), Ltryptophan-2,3,3-D_3_ (40μg/mL), (4μg/mL), L-tyrosine-^13^C_6_ (4μg/mL), caffeine-D_3_ (4μg/mL), succinic acid-2,3,3,3-D_4_ (4μg/mL), L-leucine-13C6 (4μg/mL), L-phenylalanine-^13^C_6_ (4μg/mL), N-BOC-L-tert-leucine (4μg/mL), and N-BOC-L-aspartic acid (4μg/mL) in 0.1% formic acid in water were added prior to protein precipitation. L-Leucine-D_10_, creatine-D_3_, L-tryptophan-2,3,3-D_3_, succinic acid-2,3,3,3-D_4_, and caffeine-D_3_ were purchased from CDN isotopes (Pointe-Clarie, Quebec, Canada). L-tyrosine-^13^C_6_, L-leucine-^13^C_6_, and L-phenylalanine-^13^C_6_ were purchased from Cambridge Isotope laboratories, Inc. (Tewksbury, MA). N-BOC-L-tert-leucine and N-BOC-L-aspartic acid purchased from Acros Organics (Fair Lawn, NJ.

### Sample reconstitution and monitor

Resultant dry metabolite pellet from tissue, cell, and urine were dissolved in 50 μL 0.1 % formic in water, vortexed for 10 s, sonicated for 10 minutes, and incubated for 10 min at 4 °C. The solution was centrifuged at 20,000 rcf for 10 min at 4 °C and transferred to an autosampler vial with insert. To monitor the data collection quality by UHPLC-HRMS, 50 μL of formic acid in water containing the injection standards including BOC-L-tyrosine (2 μg/mL), BOC-L-tryptophan (2 μg/mL), and BOC-D-phenylalanine (2 μg/mL) was used.

### LC-MS Analysis

Two microliters of reconstituted extract was injected onto an ACE® Excel C18-PFP, 2 μm, 2.1 × 100 mm, column (Mac-Mod Analytical, Chadds Ford, PA) and eluted at 350 μL/min (except where noted) with a gradient of Solvent A (0.1 % FA in H_2_O) and Solvent B (100% ACN): Solvent B was maintained at 0 % for 3 min, then increased to 80 % over 10 min, where it was maintained for an additional 3 min. To re-equilibrate the column, solvent B was decreased to 0 % over 0.5 min and the flow rate increased to 600 μL/min over 0.2 min; conditions were held for 3 min. The flow was then slowed to 350 μL/min over 0.2 min, and equilibrium was maintained for 2.1 min before the next injection. A Q-Exactive™ Orbitrap™ mass spectrometer (MS) (Thermo Scientific, San José, CA) in positive heated electrospray ionization (HESI) mode with Dionex UHPLC was used for all analyses. MS parameters were as follows: 3.5 kV spray voltage, 30% S-lens, 50 arb sheath gas, 15 arb auxiliary gas, 1 arb sweep gas, 275 °C gas temperature, 325 °C heated transfer tube temperature, and 35,000 mass resolution. Only positive ion data was used for comparisons in this study.

### MZmine 2 Parameters

All raw data were converted from .RAW to .mzXML format via Raw Converter.^30^ Processing was handled with MZmine v2.38.^31^ MZmine parameters were as follows: Mass Detection; MS level, 1, mass detector, centroid. Chromatogram builder; MS level, 1, minimum time span (min), 0.06, minimum height, 1.0×10^−5^, *m/z* tolerance, 0.002 *m/z* or 5.0 ppm. Smoothing; Filter width, 5. Chromatogram deconvolution; Algorithm, Local minimum search, *m/z* center calculation, MEDIAN, Remove original peak list. Isotopic peaks grouper; *m/z* tolerance, 0.002 *m/z* or 5.0 ppm, Retention time tolerance, 0.05 absolute (min), Maximum charge, 3, Representative isotope, Most intense, Remove original peak list. Join aligner, *m/z* tolerance, 0.003 *m/z* or 5.0 ppm, Weight for *m/z*, 20, Retention time tolerance, 0.05 absolute (min), Weight for RT, 20. Peak finder, Intensity tolerance, 25.0 %, *m/z* tolerance, 0.003 *m/z* or 5.0 ppm, Retention time tolerance, 0.05 absolute (min). Duplicated peak filter, Filter mode, NEW AVERAGE, *m/z* tolerance, 0.002 *m/z* or 5.0 ppm, RT tolerance, 0.05 absolute (min). Adduct search, RT tolerance, 0.05 absolute (min), *m/z* tolerance, 0.003 *m/z* or 5.0 ppm, Max relative adduct peak height, 40.0 %. Complex search, Ionization method, [M+H]^+^, Retention time tolerance, 0.05 absolute (min), *m/z* tolerance, 0.002 *m/z* or 5.0 ppm, Max complex peak height, 50.0 %.

### XCMS Parameters

The same .mzXML files for cells, tissues, and urine analyzed by MZMine were uploaded to the XCMS cloud via the Scripps XCMS webpage (https://xcmsonline.scripps.edu).

XCMS parameters were as follows: General; Name, UPLC/Q-Exactive, Comment, UPLC-QExactive, Polarity, positive, Retention time format, minutes. Feature Detection; ppm, 5, minimum peak width, 5, maximum peak width, 20, Signal/Noise threshold, 4, mzdiff, 0.01, Integration method, 1, prefilter peaks, 3, prefilter intensity, 100, Noise Filter, 0. Retention Time Correction; profStep, 1. Alignment; mzwid, 0.025, bw, 5, minfrac, 0.5, minsamp, 1, max, 100. Statistics; Statistical Test, NA, p-value threshold (highly significant features), 0.01, fold change threshold (highly significant features), 1.5, p-value threshold (significant features), 0.05, value, into. Annotation; Search for, isotopes + adducts, ppm, 5, m/z absolute error, 0.015. Identification; adducts, [M+H]^+^ [M+NH4]^+^ [M+Na]^+^ [M+H-H2O]^+^ [M+K]^+^ [M+2Na-H]+ [M+2H]^2+^, ppm, 15, sample biosource, NA, pathway ppm deviation, 5, significant list p-value cutoff, AUTO. Visualization; EIC width, 100.

### Raw peak picking parameters

Raw spectral peak intensity data were extracted manually and then processed using MetaboAnalyst 4.0 based on a mass tolerance of ±0.005 ppm and retention time tolerance 30.^32^ In the case of zero intensity results, gap filling was performed to replace zero values (using half the minimum value) only when the feature was present in 80% of the samples.^29^ The data were filtered by blank feature filtration (BFF) to remove features with ≥10% signal contribution from the background. The data was log 2 transformed and auto scaled.^12^ Metabolites were annotated based on *m/z* exact mass (molecular weight tolerance of 5 ppm) matching to the HMDB database (https://hmdb.ca/spectra/ms/search).

### Blank feature filtering to eliminate noise

The untargeted metabolomics workflow run blanks at systematic intervals between randomized blocks of experimental data. Blank feature filtering (BFF) approach was used to remove features not originating from the sample. For each detected metabolite, a signal-to-noise metric was calculated by comparing sample peak heights to blank peak heights using the following equation:

**BFF=Q1(S1**…**Sn)−(Ave (B1**…**Bn) + c x SD(B1**…**Bn))**

**BFF = Q-LLQ**

BFF = Blank feature filtering

B1…Bn = Blank sample 1 to n

S1…Sn = Experimental sample 1 to n

Ave = Average

SD = Standard deviation

Q1 = Quartile 1

Q = Experimental Signal

LLQ = Noise signal

LLQ = (Ave(B1…Bn)+cxSD(B1…Bn))

Q1-LLQ = Sigal-to-Noise discrimination signal

### Chemometrics Analysis

MetaboAnalyst 6.0 was used for data processing with the following parameters: peak intensity table, samples in columns unpaired, missing value estimation used to replace by a small value (half of the minimum positive value in the original data, none of the features were removed in this step), data filtering by relative standard deviation (RSD = SD/mean), normalized by sum (to correct the instrumental and the technical variation), data transformed using log transformation, and data scaled using autoscaled (to allow a more direct comparison between features of greatly varying intensities). Evaluation of different multivariate statistical analysis models including principal component analysis (PCA), partial least square discrimination analysis (PLS-DA), orthogonal-partial least square discrimination analysis (OPLS-DA) and random forest-based variable importance in projection (VIP) analysis were conducted using MetaboAnalyst 5.0.^32^ Metabolic pathway analysis was conducted using the Kyoto Encyclopedia of Genes and Genomes (KEGG) pathway database by matching metabolite sets with the human metabolome (https://www.genome.jp/kegg/pathway.html). Metabolite set enrichment (fold enrichment) were investigated using MetaboAnalyst 6.0 (open source R package)^33^.

## RESULTS and DISCUSSION

UHPLC-HRMS-based untargeted metabolomics data were acquired from cells, tissues, and urine by reversed-phase UHPLC in positive ionization mode scanning from *m/z* 70 to 1000 at 35,000 mass resolution. Raw data were processed with MZmine 2 and XCMS after conversion from .RAW to .mzXML format with MSConvert. For simplicity, only positive ion data were analyzed. Peak-picking was compared first between MZmine and XCMS. Secondly, the impact of software choice on metabolic profiling results was evaluated via different chemometric tests (PCA and PLS-DA). Finally, differences between MZmine and XCMS in biomarker prediction and identification were investigated.

### Metabolic peak picking differs between MZmine 2 and XCMS

Cells, tissues, and biofluids are all commonly tested samples in cancer metabolomics. Peak-picking was evaluated for each sample set and compared between MZmine 2 and XCMS (**Figure 1**). All data were filtered using BFF as mentioned previously.^12^ As shown in Figure 1A, the total number of metabolic peaks identified greatly differed between MZmine 2 and XCMS, regardless of sample type. MZmine 2 identified ∼25K peaks in urine, ∼5K peaks in cells, and ∼1.6K peaks in tissue, whereas XCMS identified ∼11K in urine, ∼2K in cells, and ∼1.4K in tissue, respectively (**Figure 1**).

**Figure 1.**
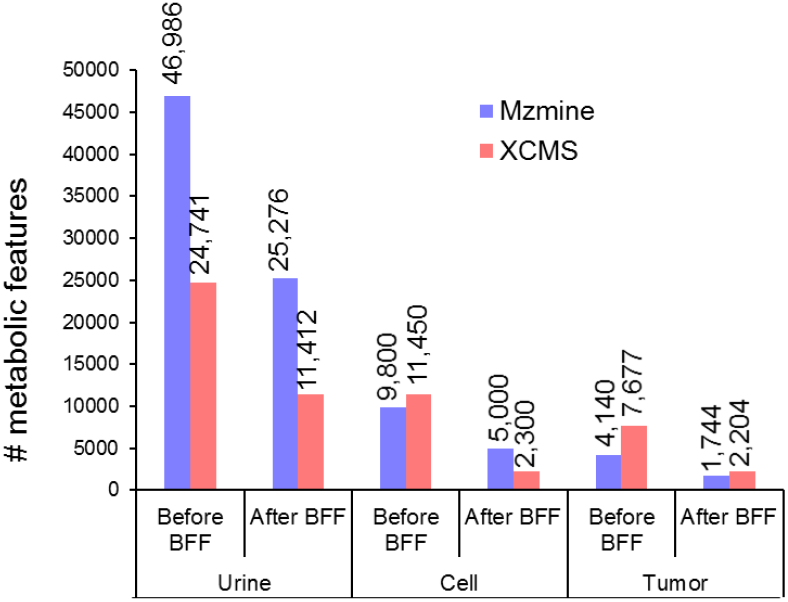
Metabolic peaks between MZmine and XCMS before and after blank feature filtering. Shown are the number of identified peaks in urine, cell, and tissue before and after Blank Feature Filtering (BFF) by MZmine and XCMS.

Recent studies have noted MZmine and XCMS perform peak detections differently^9,16^ and differ in component detections by ∼10% for UHPLC-HRMS and ∼20% for GC-MS ^14^. Similarity between MZmine and XCMS-generated data sets were evaluated by looking at the overlap (0.001 *m/z*, 0.1 min, and 0.01 min retention time) in feature lists for both software packages. Large discrepancies were observed in all sample types (**Figure 2A-D**). We first observed the peak discrepancy across the sample types using two different retention time (RT) tolerances: 0.1 min and 0.01 min. Using a RT of 0.1 min, we identified 38.7 % (10,733 peaks), 20% (1468 peaks), and 31% (724 peaks) of the intersected peaks in urine, cell, and tissue samples, respectively, by both XCMS and MZmine (Figure 2A). Interestingly, the 0.01 min RT window demonstrated 0%, 1.2% (106 peaks), and 3% (81 peaks) intersection for urine, cells, and tissue samples, respectively (**Figure 2B**). These observations indicate that the choice of RT tolerance can have a significant influence on the identification and comparison of peaks across different sample types by different peak picking tools in metabolomics analysis.

**Figure 2.**
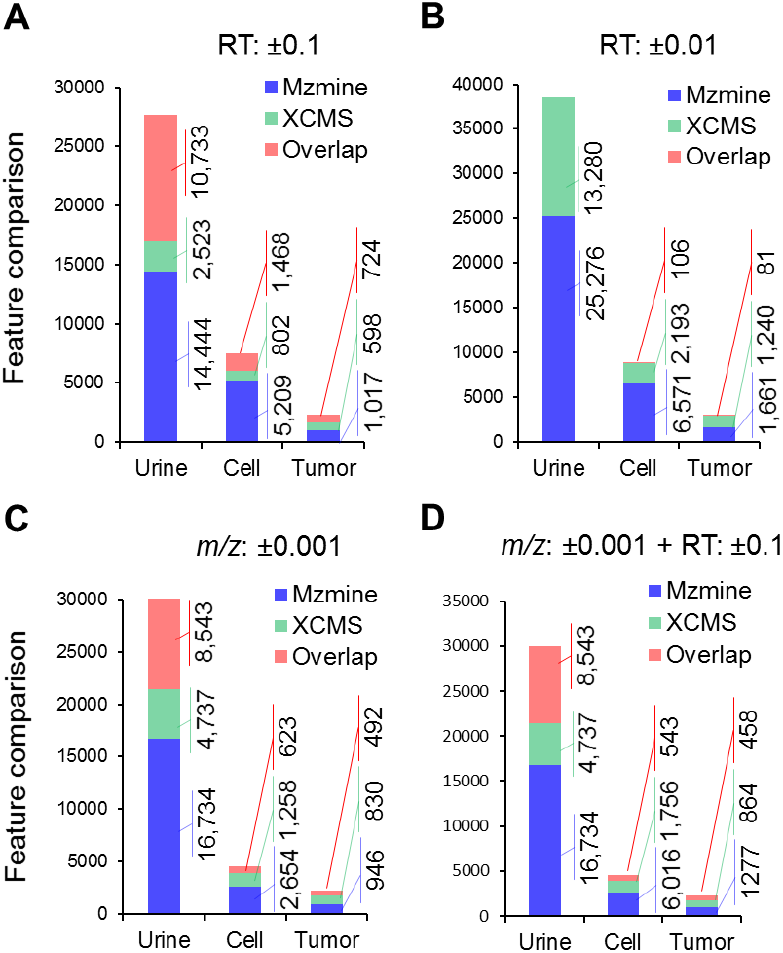
Comparative metabolic peak picking by MZmine and XCMS. Stacked bar graph showing unique and overlapping peaks found in MZmine and XCMS at 0.1 (A) and 0.01 (B) RT tolerance, respectively, as well as 0.001 *m/z* (C), and 0.001 *m/z* ±0.1 RT window together (D), respectively.

Furthermore, we evaluated the peak intersection across the sample types using a *m/z* tolerance of 0.001, standard tolerance for high resolution MS-based metabolomics, and identified that the *m/z* dimension of 3 ppm reduced peak coherence across the data matrices, where 28.5% (8,543 peaks), 13.7% (623 peaks), and 21.7% (492 peaks) overlapped peaks were identified in urine, cells, and tissue samples, respectively (**Figure 2C**). In metabolomics, the combination of precursor *m/z* and RT are used to derive structural information of a metabolite and provide data for statistical anaylsis.^34^ Hence, we explored the peak comparison between MZmine 2 and XCMS based on both *m/z* and RT (**Figure 2D**). In urine samples (healthy vs cancer), we observed 28.5% (8,543 peaks) overlapping peaks between MZmine and XCMS with 0.001 *m/z* and 0.1 RT together (**Figure 2D**). In case of cells (normal vs cancer), we observed 6.5% (543 peaks) overlapped peaks between MZmine 2 and XCMS with 0.001 *m/z* and 0.1 RT together (**Figure 2D**). In the case of tissues (normal vs tumor), we observed 17.6% (458 peaks) overlapped peaks between MZmine and XCMS with 0.001 *m/z* and 0.1 RT together (**Figure 2D**). Collectively, our data analysis suggests low coherence regarding peak picking between MZmine and XCMS software regardless of the sample matrix.

To understand the detailed sources of peak picking discrepancies, we further dissected the source of metabolic feature discrepancies throughout the total ion chromatogram (TIC) by looking at different solvent gradient portions of the chromatographic run, including the isocratic portion (0% A from 0-3 min), the gradient portion (3-13 min), and isocratic elution (80% B from 13-16 min) (**Figure 3A&B**). Interestingly, we observed that under the gradient part of the separation, more features were identified in the case of urine and cells, in contrast for tissue samples, the early isocratic part of the run (0-3 min) contributed the most features **(Figure 3C)**. This could be due to the lower overall signal or may be reflective of more polar analytes present in the tissues tested since the early isocratic portion is 0% acetonitrile. This would need to be investigated further. We found that the 80% isocratic part from 13-16 min contributed minimally to the total metabolic features identified (**Figure 3C**). This discrepancy suggests that the choice of solvent gradient and the time duration of different portions of the run may impact the detection and identification of metabolomic features.

**Figure 3.**
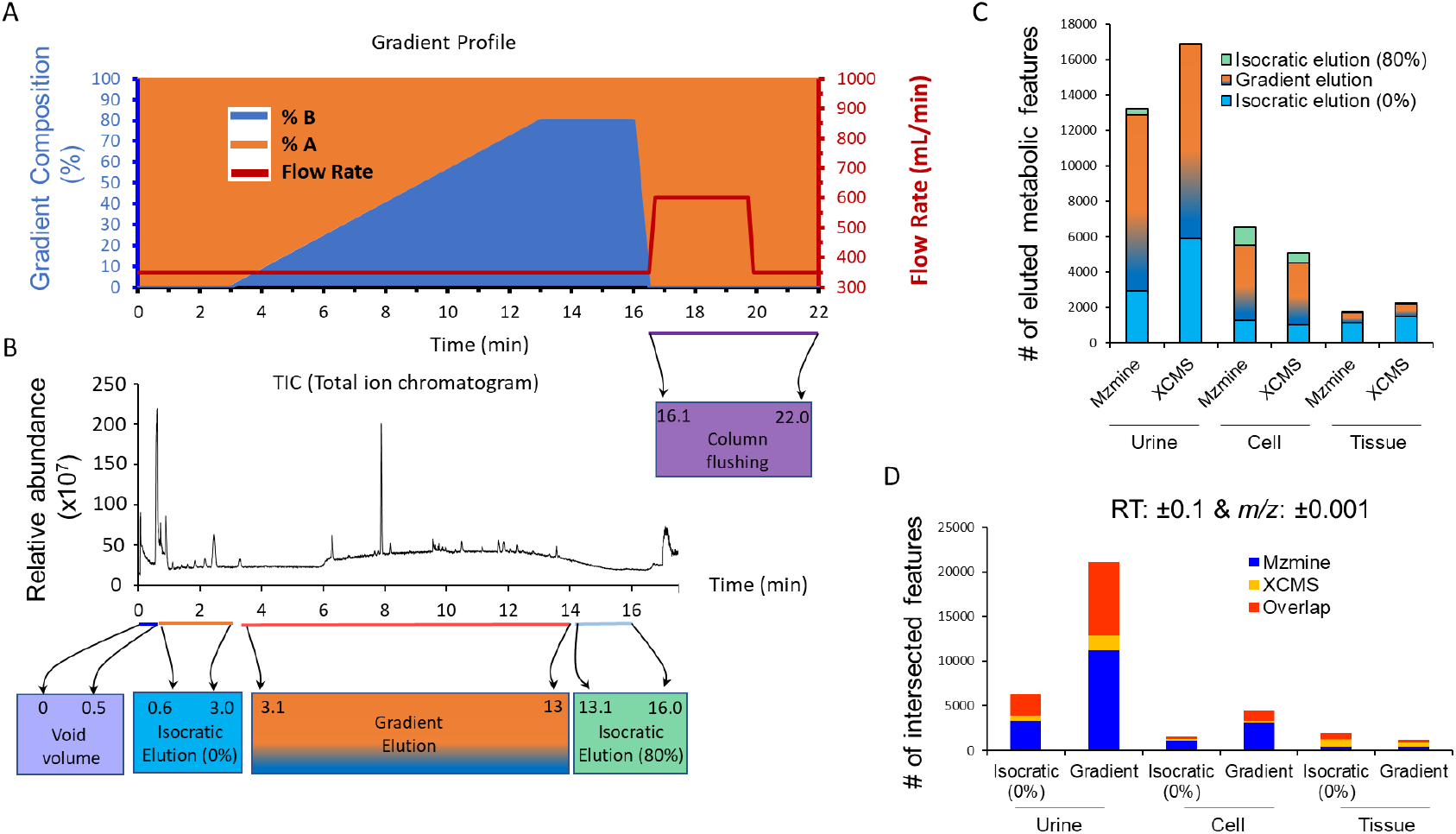
Impact of different solvent gradient compositions in metabolic peak picking analysis. A typical solvent gradient profile of untargeted metabolomics analysis using UHPLC-HRMS (A). Total ion chromatogram (TIC) with indicated solvent elution phases such as isocratic elution at 0% (0.6-3.0 min), gradient elution (3.1-13.0 min), and isocratic elution at 80% (13.1-16.0 min) at the bottom (B). Stacked bar graph shows number of eluted metabolic features identified by MZmine and XCMS for urine, cell, and tissue specimens, respectively (C). Number of intersected features between MZmine and XCMS at combined 0.1 RT and 0.001 *m/z* (D).

Next, we wanted to understand whether different solvent gradient aspects of a separation were involved in discrepancies of metabolic peak picking by different tools. MZmine and XCMS were employed to investigate overlapping metabolic features generated by the initial isocratic elution (0-3 min) and gradient elution (3-13 min). In Figure 2, it was demonstrated that using a combination of 0.1 RT and 0.001 m/z tolerance yielded better peak overlap between MZmine and XCMS tools. Therefore, here we only demonstrated combination of 0.1 min RT and 0.001 *m/z* tolerance (Figure 3D). We observed overlapping peaks of 37.4% and 39.3% in urine, 15.3% and 25.1% in cells, and 36.1% and 18% in tissue for the initial isocratic elution (0-3 min) and gradient elution (3-13 min), respectively (**Figure 3D**). Here, we observed that the most overlapped peaks were from the isocratic elution phase (0-3 min) phase and the quantity of intersected peaks slightly dropped in the gradient phase except for cell samples. Overall, regardless of sample types, different solvent gradient phases, different RT, or *m/z* tolerances, we observed low coherence or high discrepancies in metabolic peak picking between the MZmine and XCMS tools.

### Global metabolic landscape differs between MZmine and XCMS peak picking

To evaluate the global metabolic landscape between MZmine and XCMS peak picking, different chemometric approaches including PCA and hierarchical clustering were utilized.^35–37^

Despite the high level of discrepancies in peak picking, MZmine and XCMS produced relatively similar profiles in principal component analysis (PCA) in the case of healthy versus prostate cancer urine (**Figure S1**). Also, a similar trend of metabolic profiles was identified between MZmine and XCMS peaks for normal versus tumor cells (**Figure S2**) and normal versus tumor tissue (**Figure S3**) with PCA. However, hierarchical clustering heatmap analysis of the top 40 metabolic features in the case of normal (n=3) versus tumor tissues (n=3) were contrary to PCA analysis with marked differences observed between the XCMS and MZmine data sets (**Figure 4A&B**). Likewise, the top 40 metabolic features identified between MZmine and XCMS peak lists were also found to be very different for normal (n=3) versus oral cancer cells (n=3) (**Figure 4C&D**) and normal (n=10) versus prostate cancer urine specimens (n=10) (**Figure 4E&F**). This finding is not surprising, however, considering that heatmaps are directly affected by feature identity, whereas chemometric analyses are broader in scope. Another group has noted similar results when comparing hierarchical clustering and multivariate statistical tests among Haystack, MZmine, and manual analysis.^38^ Results herein described are largely in agreement with the community consensus that individual feature discrepancies among different peak-picking packages are largely irrelevant to broader conclusions in global metabolomics. Further investigation into how these differences can affect biomarker identification and prediction is warranted.

**Figure 4.**
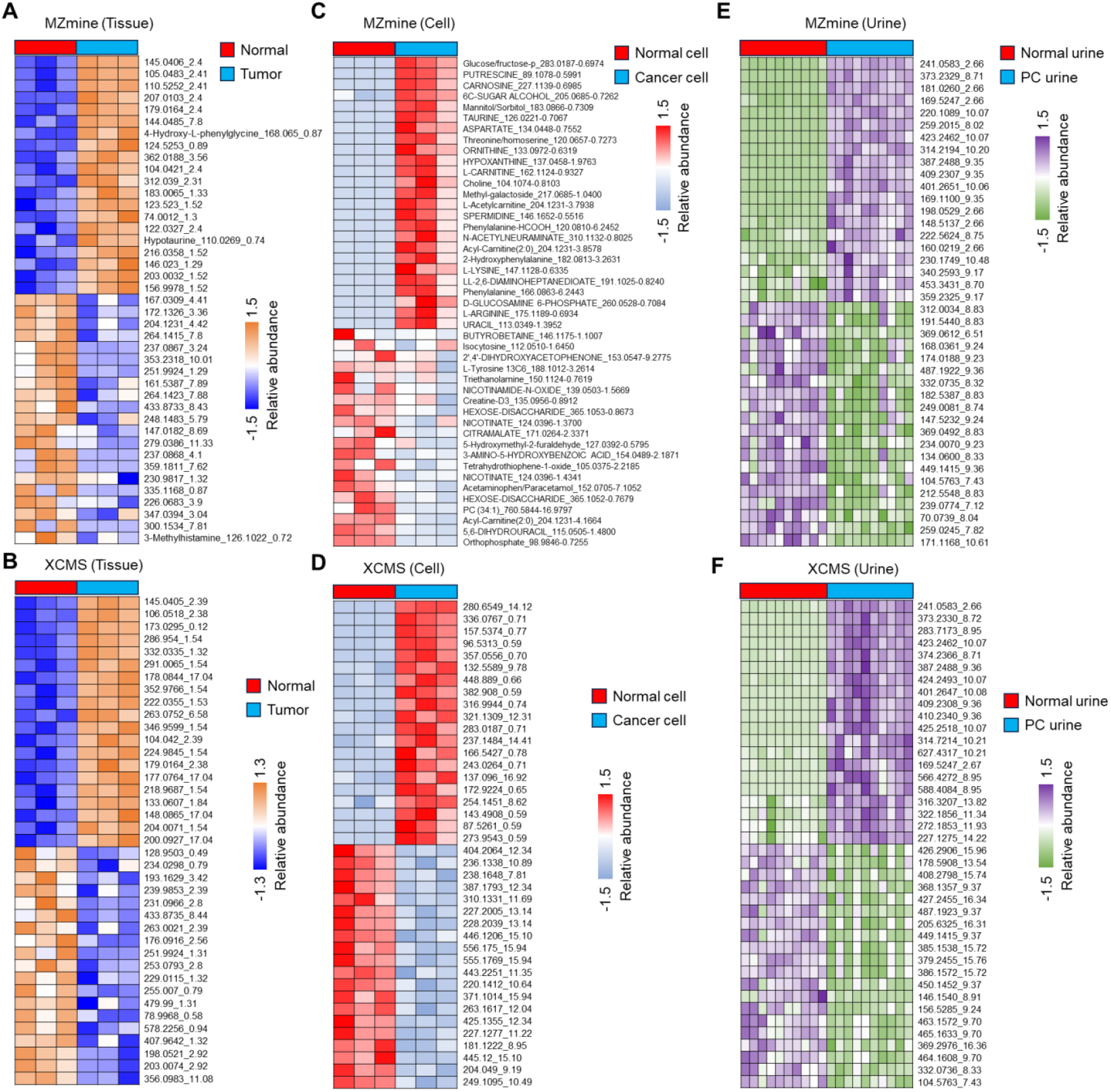
Metabolic profile differences between normal and different disease states between MZmine and XCMS. Shows a heatmap of top 40 metabolic feature (m/z_RT) abundances in normal tissue (n=3) vs tumor tissue (n=3) samples by MZmine (A) and XCMS (B). Shows a heatmap of top 40 metabolic feature abundances in normal cell (n=3) vs cancer cell (n=3) samples by MZmine (C) and XCMS (D). Shows a heatmap of top 40 metabolic feature abundances in normal urine (n=10) vs PC urine (n=10) samples by MZmine (E) and XCMS (F), respectively. Relative abundance shown by z-score calculation.

### Biomarker discrepancies between MZmine and XCMS-based peak picking, regardless of sample matrix

Metabolomic profiling using MZmine and XCMS revealed distinct variations in biomarker detection across multiple biological matrices. ROC curve analysis demonstrated robust diagnostic capabilities for both platforms (**Figure S4**), with AUC values ranging from 0.66 to 1.0. While MZmine exhibited marginally superior performance in tissue analysis (AUC = 0.86) compared to XCMS (AUC = 0.66) when utilizing the top 5 metabolites, both platforms showed comparable efficacy when analyzing larger feature sets (n > 25) (**Figure S4**). This convergence in performance suggests equivalent analytical robustness for comprehensive biomarker discovery applications.

Differential metabolite analysis via volcano plots (**Figure S5**) employed stringent filtering criteria (p < 0.05, FC ≥ 1.5) across tumor tissue, cancer cell, and urine specimens. Statistical significance analysis revealed distinct metabolic signatures between normal and tumor samples, with both platforms identifying unique feature sets. The comparative analysis highlighted platform-specific differences in peak detection and integration algorithms, particularly evident in the distribution of significantly altered metabolites (designated by red for upregulation and blue for downregulation) (**Figure S5**).

Further assessment of the top 10 significant metabolic features (**Figure 5**) based on m/z_RT values demonstrated notable discrepancies between MZmine and XCMS processing workflows. Feature detection patterns varied substantially across sample matrices, with distinct differences observed in thyroid tumor tissue, oral cancer cells, and PC urine samples (**Figure 5**). Statistical validation (p = 0.01-0.04) and fold-change magnitude visualization revealed platform-dependent variations in metabolite identification and quantification, underscoring the importance of careful platform selection and validation in metabolomics investigations.

**Figure 5.**
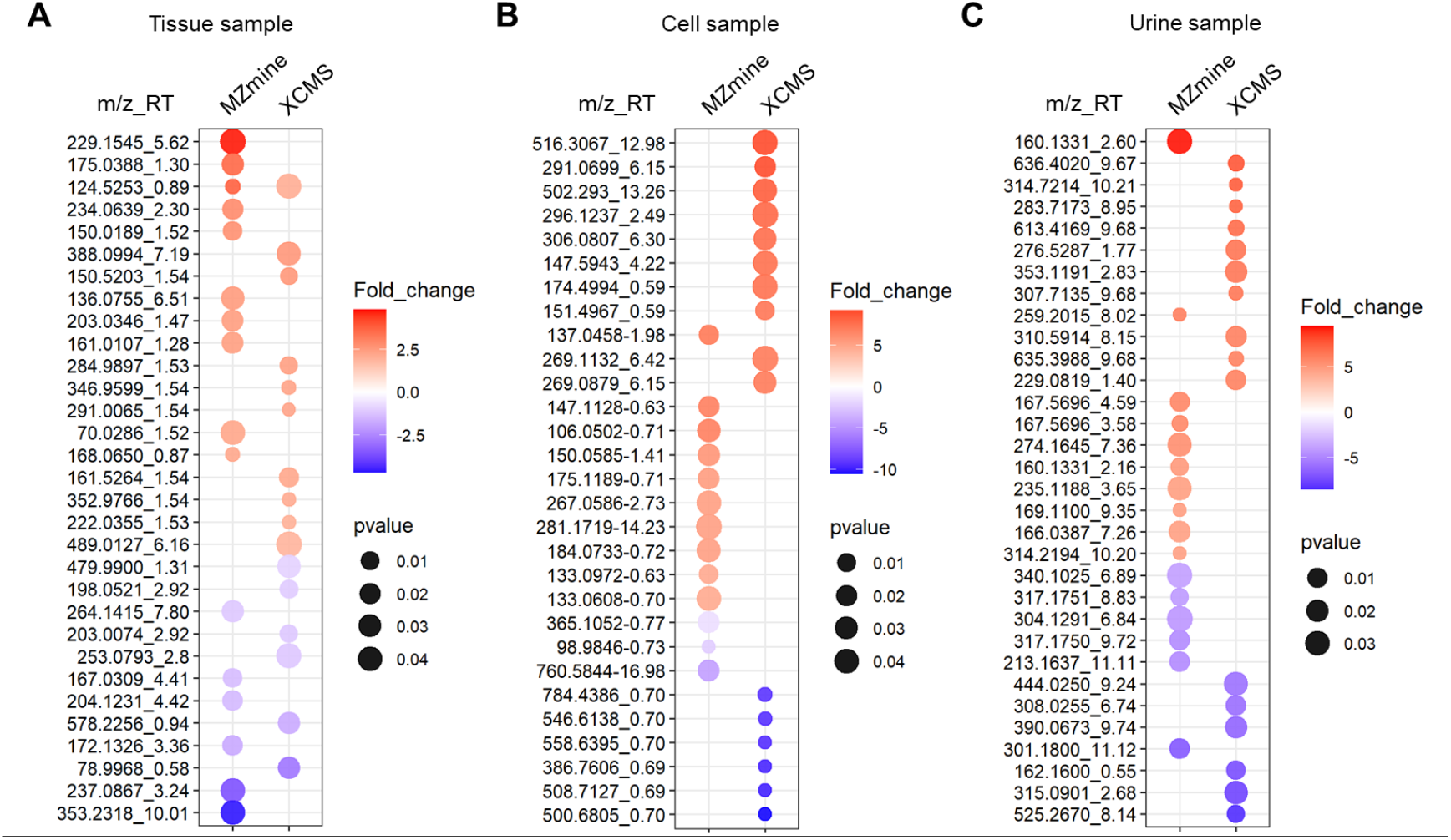
Distinct discrepancies in significant metabolic feature analyzed by MZmine and XCMS. The dot plot shows the top 10 significant metabolites based on m/z_RT values in tissue (normal vs thyroid tumor) (A), cell (normal vs oral cancer cells) (B), and urine (normal vs PC urine samples) (C). Red dots indicate upregulated metabolites while blue dots indicate downregulated ones. Dot size corresponds to p-value significance (0.01-0.04). The intensity of color represents fold change magnitude, with darker shades indicating greater changes.

**Figure 6.**
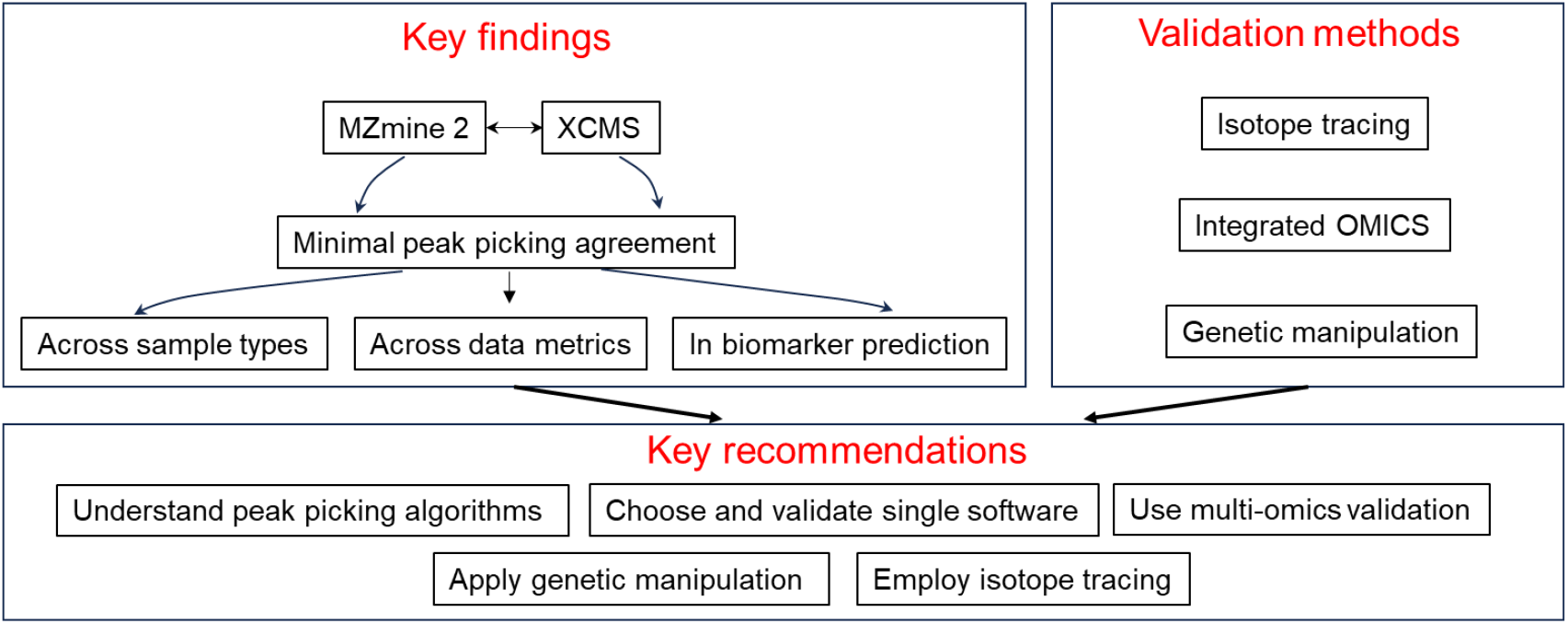
Summary on non-targeted metabolic biomarker analysis and validation.

Moreover, to better understand the accuracy between MZmine and XCMS for untargeted metabolomics-based biomarker analysis, we conducted manual peak picking as mentioned previously^39^ and compared manual peak list with MZmine or XCMS peak (**Figure S6**). Interestingly, we did not find a significant peak intersection between manual and MZmine or XCMS (**Figure S6**). However, MZmine showed improved coherence with the manual peak list compared to XCMS. Our findings underscore the critical need for standardization of metabolomics data processing algorithms, suggesting that refinement of automated peakpicking methods could significantly enhance the reliability and reproducibility of biomarker discovery in untargeted metabolomics studies.

### Summary

Our comprehensive investigation of untargeted metabolomics data processing revealed fundamental discrepancies between MZmine and XCMS platforms across diverse biological matrices. Analysis of cells, tissues, and biofluids demonstrated minimal agreement between these software tools in peak detection, regardless of sample types or analytical conditions. Both platforms showed notably low coherence in biomarker prediction and reliability analysis, with differences particularly evident when evaluating peak detection across various chromatographic conditions, retention times, and m/z tolerances. This systematic comparison highlighted significant variability in feature detection during different solvent gradient phases and demonstrated the critical impact of software choice on downstream metabolomics analyses.

Based on these findings, we propose a hierarchical validation framework for metabolomics investigations. The framework begins with fundamental algorithm understanding and proper software implementation, followed by rigorous platform selection and validation protocols. This foundation should be supplemented with multi-omics integration, incorporating upstream genomics, transcriptomics, and proteomics data to validate metabolomics findings. Further validation through genetic manipulation and isotope tracing studies can provide additional confirmation of biomarker responses and metabolic pathway alterations. This comprehensive approach addresses the current limitations in metabolomics data processing while establishing a robust pipeline for reliable biomarker discovery and validation.

## ASSOCIATED CONTENT

## Supporting Information

The Supporting Information is available free of charge on the ACS Publications website.

**Figure S1**. Global metabolic profile analysis of normal versus prostate cancer urine metabolic features based on peak lists generated by MZmine and XCMS, respectively.

**Figure S2**. Global metabolic profile analysis of normal versus oral cancer cell metabolic features based on peak lists generated by MZmine and XCMS, respectively.

**Figure S3**. Global metabolic profile analysis of normal versus thyroid tumor tissue metabolic features based on peak lists generated by MZmine and XCMS, respectively.

**Figure S4**. ROC (Receiver Operating Characteristic) curves comparing the diagnostic performance of metabolic features identified by MZmine (top row) and XCMS (bottom row) across three sample types: tissue, urine, and cell samples.

**Figure S5**. Biomarker discrepancy analysis using volcano plot analysis.

**Figure S6**. Intersection among manual peak list, MZmine, and XCMS peak list from PC urine versus healthy urine specimens.

## AUTHOR INFORMATION

### Author Contributions

I.M. conceptualization, data collection, data analysis, and wrote the manuscript. T.A.H data collection, data analysis and review the manuscript. L.E.M. review. T.J.G. Supervision, review and editing the manuscript.

## ACKNOWLEDGMENT

This work was funded by the National Institutes of Health grant U24DK097209.

**Notes**

The author declares no competing interest.

## Supplementary Figures

**Figure S1.**
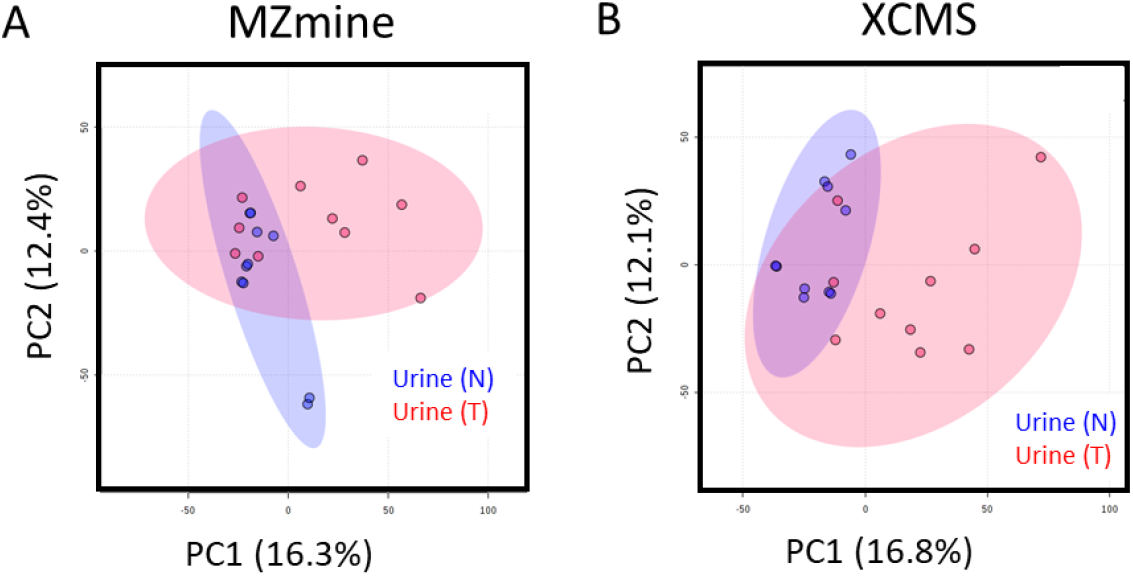
Global metabolic profile analysis of normal (N) versus prostate cancer (T) urine metabolic features based on peak lists generated by MZmine and XCMS, respectively.

**Figure S2.**
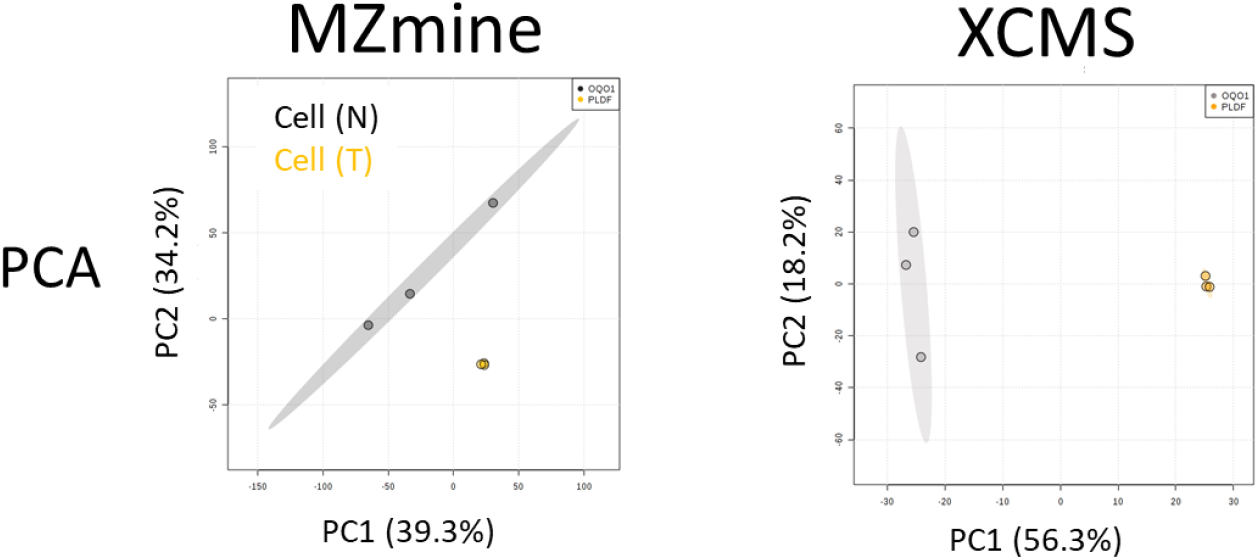
Global metabolic profile analysis of normal (N) versus oral cancer cell (T) metabolic features based on peak lists generated by MZmine and XCMS, respectively.

**Figure S3.**
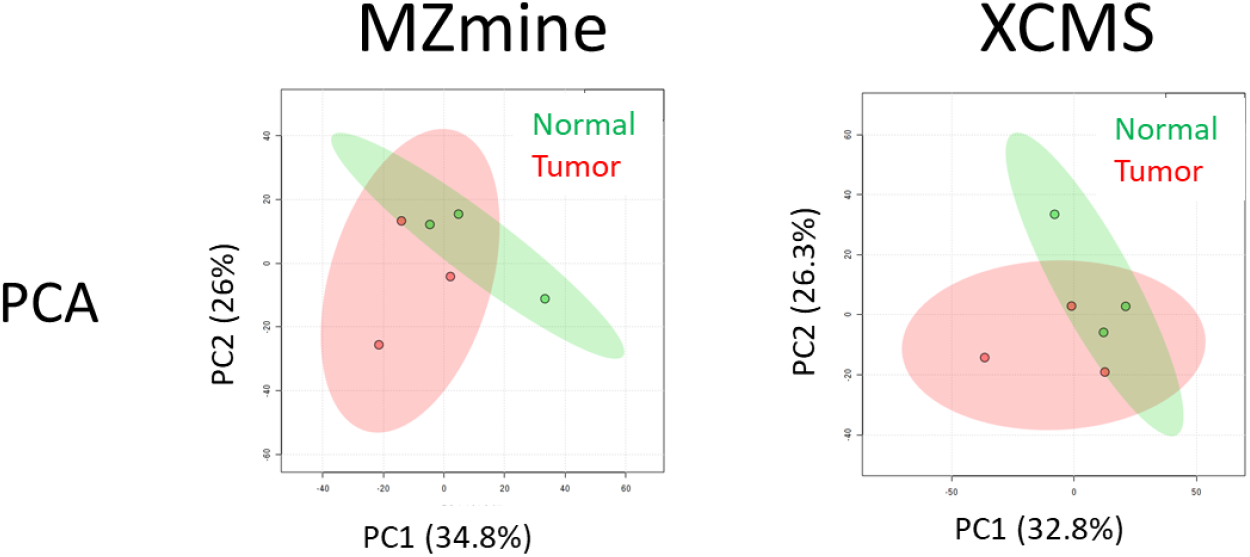
Global metabolic profile analysis of normal (N) versus thyroid tumor tissue (T) metabolic features based on peak lists generated by MZmine and XCMS, respectively.

**Figure S4.**
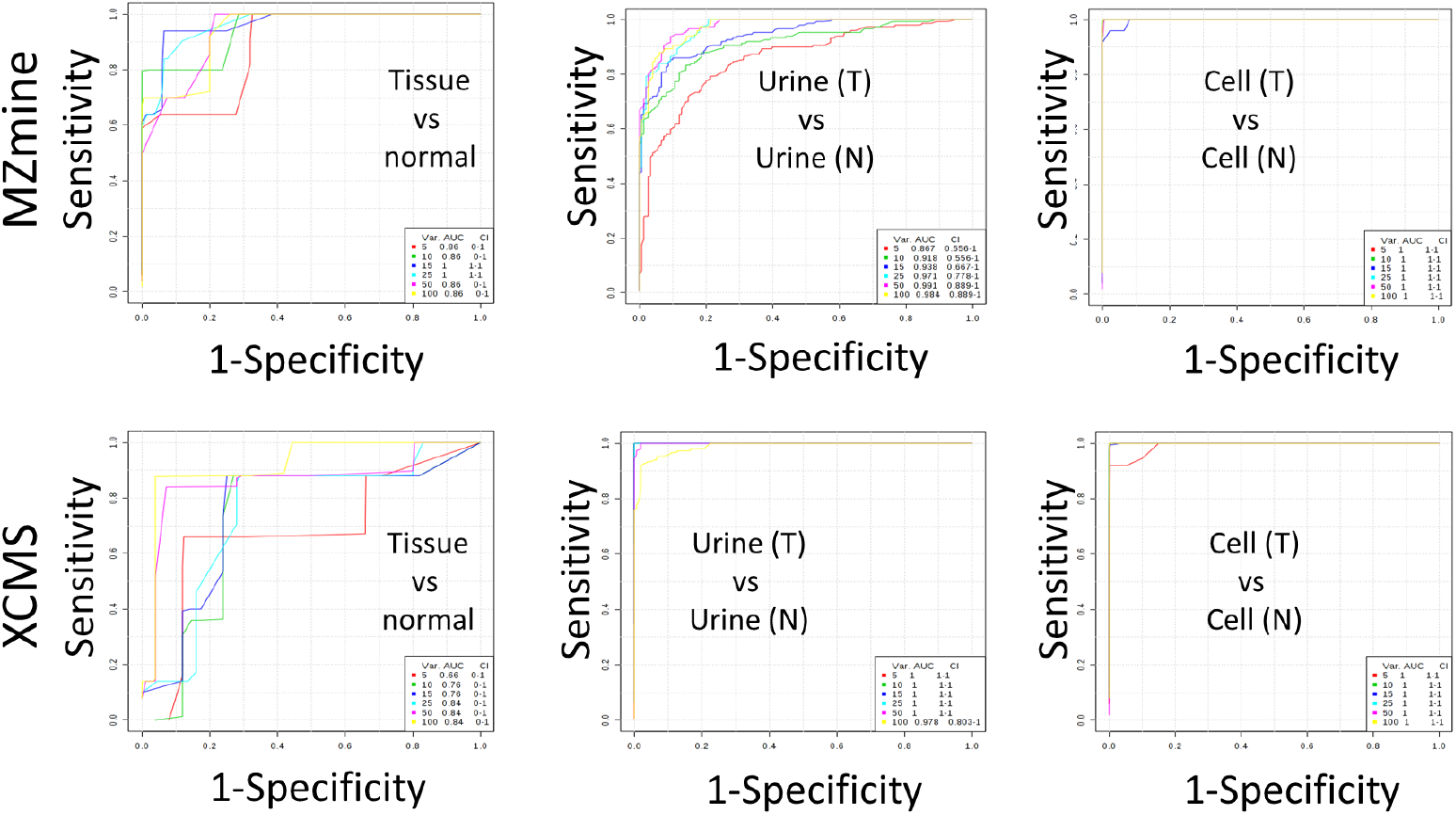
ROC (Receiver Operating Characteristic) curves comparing the diagnostic performance of metabolic features identified by MZmine (top row) and XCMS (bottom row) across three sample types: tissue, urine, and cell samples. Each curve plots sensitivity versus 1-specificity, with different colored lines representing distinct metabolic features. The accompanying values show AUC (Area Under Curve) and CI (Confidence Interval) for each feature. The analysis compares tumor (T) versus normal (N) samples for each specimen type.

**Figure S5.**
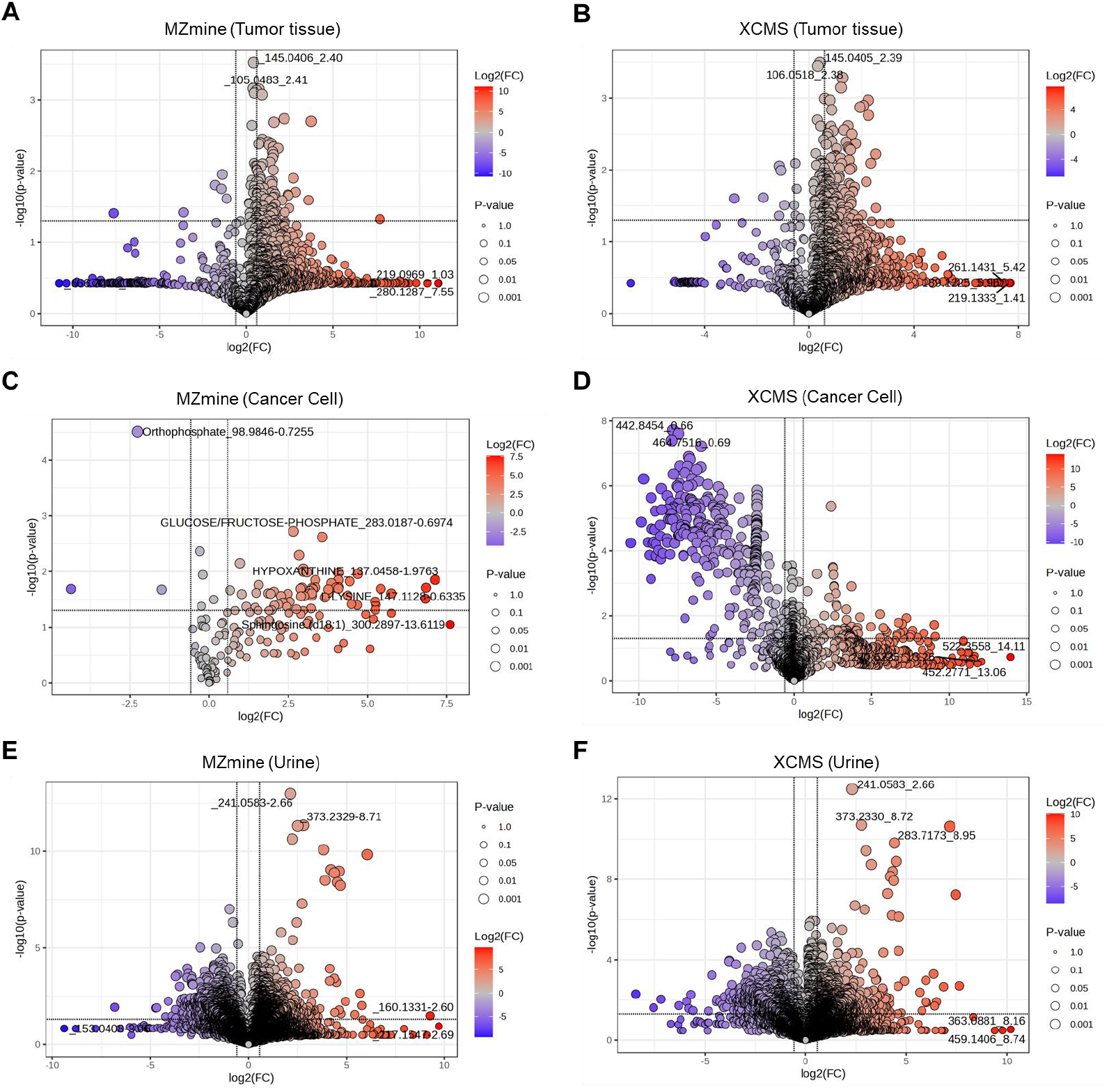
Biomarker discrepancy analysis using volcano plot analysis. Comparisons of metabolite levels between normal and tumor samples across three different types of specimens such as tumor tissue (A & B), cancer cell (C & D), and urine (E&F) analyzed using two metabolomics software packages (MZmine and XCMS). Each plot displays log2 fold change in x-axis with threshold ≥ ±1.5 versus -log10 p-value (y-axis) with p-value threshold <0.05. Significantly altered metabolites are indicated by colored dots (red: upregulated, blue: downregulated), with dot size corresponding to statistical significance (p-value thresholds: 0.001, 0.01, 0.05, 0.1).

**Figure S6.**
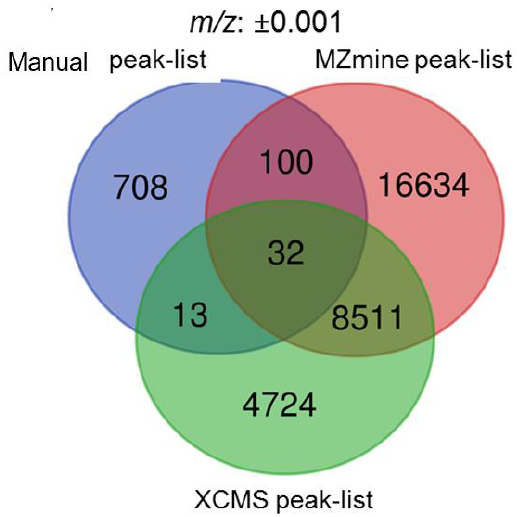
Intersection among manual peak list, MZmine, and XCMS peak list from PC urine versus healthy urine specimens. Venn diagram showing unique and overlap peaks found among raw peak list, MZmine and XCMS peak list at 0.001 *m/z* window.

## REFERENCES

(1) Schrimpe-Rutledge, A. C.; Codreanu, S. G.; Sherrod, S. D.; McLean, J. A. Untargeted Metabolomics Strategies – Challenges and Emerging Directions. J. Am. Soc. Mass Spectrom. 2016, 27 (12), 1897–1905. 10.1007/s13361016-1469-y.

(2) Gertsman, I.; Barshop, B. A. Promises and Pitfalls of Untargeted Metabolomics. J. Inherit. Metab. Dis. 2018, 41 (3), 355–366. 10.1007/s10545-017-0130-7.

(3) O’Kell, A. L.; Garrett, T. J.; Wasserfall, C.; Atkinson, M. A. Untargeted Metabolomic Analysis in Naturally Occurring Canine Diabetes Mellitus Identifies Similarities to Human Type 1 Diabetes. Sci. Rep. 2017, 7 (1), 9467. 10.1038/s41598-017-09908-5.

(4) Menzies, V.; Starkweather, A.; Yao, Y.; Kelly, D. L.; Garrett, T. J.; Yang, G.; Booker, S.; Swift-Scanlan, T.; Mahmud, I.; Lyon, D. E. Exploring Associations Between Metabolites and Symptoms of Fatigue, Depression and Pain in Women With Fibromyalgia: Biol. Res. Nurs. 2020. 10.1177/1099800420941109.

(5) Lynch Kelly, D.; Farhadfar, N.; Starkweather, A.; Garrett, T. J.; Yao, Y.; Wingard, J. R.; Mahmud, I.; Menzies, V.; Patel, P.; Alabasi, K. M.; Lyon, D. Global Metabolomics in Allogeneic Hematopoietic Cell Transplantation Recipients Discordant for Chronic Graft-versus-Host Disease. Biol. Blood Marrow Transplant. 2020. 10.1016/j.bbmt.2020.06.014.

(6) Lee, B.; Mahmud, I.; Marchica, J.; Dereziński, P.; Qi, F.; Wang, F.; Joshi, P.; Valerio, F.; Rivera, I.; Patel, V.; Pavlovich, C. P.; Garrett, T. J.; Schroth, G. P.; Sun, Y.; Perera, R. J. Integrated RNA and Metabolite Profiling of Urine Liquid Biopsies for Prostate Cancer Biomarker Discovery. Sci. Rep. 2020, 10 (1), 1–17. 10.1038/s41598-020-60616-z.

(7) Mahieu, N. G.; Patti, G. J. Systems-Level Annotation of a Metabolomics Data Set Reduces 25 000 Features to Fewer than 1000 Unique Metabolites. Anal. Chem. 2017, 89 (19), 10397–10406. 10.1021/acs.anal-chem.7b02380.

(8) Mahmud, I.; Sternberg, S.; Williams, M.; Garrett, T. J. Comparison of Global Metabolite Extraction Strategies for Soybeans Using UHPLC-HRMS. Anal. Bioanal. Chem. 2017, 409 (26), 6173–6180. 10.1007/s00216-017-0557-6.

(9) Myers, O. D.; Sumner, S. J.; Li, S.; Barnes, S.; Du, X. Detailed Investigation and Comparison of the XCMS and MZmine 2 Chromatogram Construction and Chromatographic Peak Detection Methods for Preprocessing Mass Spectrometry Metabolomics Data. Anal. Chem. 2017, 89 (17), 8689–8695. 10.1021/acs.analchem.7b01069.

(10) Alonso, A.; Marsal, S.; Julià, A. Analytical Methods in Untargeted Metabolomics: State of the Art in 2015. Front. Bioeng. Biotechnol. 2015, 3. 10.3389/fbioe.2015.00023.

(11) Riekeberg, E.; Powers, R. New Frontiers in Metabolomics: From Measurement to Insight. F1000Research 2017, 6. 10.12688/f1000research.11495.1.

(12) Patterson, R. E.; Kirpich, A. S.; Koelmel, J. P.; Kalavalapalli, S.; Morse, A. M.; Cusi, K.; Sunny, N. E.; McIntyre, L. M.; Garrett, T. J.; Yost, R. A. Improved Experimental Data Processing for UHPLC–HRMS/MS Lipidomics Applied to Nonalcoholic Fatty Liver Disease. Metabolomics 2017, 13 (11), 142. 10.1007/s11306-017-1280-1.

(13) Wang, L.; Xing, X.; Chen, L.; Yang, L.; Su, X.; Rabitz, H.; Lu, W.; Rabinowitz, J. D. Peak Annotation and Verification Engine for Untargeted LC-MS Metabolomics. Anal. Chem. 2019, 91 (3), 1838–1846. 10.1021/acs.analchem.8b03132.

(14) Coble, J. B.; Fraga, C. G. Comparative Evaluation of Preprocessing Freeware on Chromatography/Mass Spectrometry Data for Signature Discovery. J. Chromatogr. A 2014, 1358, 155–164. 10.1016/j.chroma.2014.06.100.

(15) Rafiei, A.; Sleno, L. Comparison of Peak-Picking Workflows for Untargeted Liquid Chromatography/High-Resolution Mass Spectrometry Metabolomics Data Analysis. Rapid Commun. Mass Spectrom. 2015, 29 (1), 119–127. 10.1002/rcm.7094.

(16) Hohrenk, L. L.; Itzel, F.; Baetz, N.; Tuerk, J.; Vosough, M.; Schmidt, T. C. Comparison of Software Tools for LC-HRMS Data Processing in Non-Target Screening of Environmental Samples. Anal. Chem. 2019, acs.analchem.9b04095. 10.1021/acs.analchem.9b04095.

(17) Kern, S. E. Why Your New Cancer Biomarker May Never Work: Recurrent Patterns and Remarkable Diversity in Biomarker Failures. Cancer Res. 2012, 72 (23), 6097–6101. 10.1158/0008-5472.CAN-12-3232.

(18) Ioannidis, J. P. A. Biomarker Failures. Clin. Chem. 2013, 59 (1), 202–204. 10.1373/clinchem.2012.185801.

(19) Mahmud, I.; Tian, G.; Wang, J.; Hutchinson, T. E.; Kim, B. J.; Awasthee, N.; Hale, S.; Meng, C.; Moore, A.; Zhao, L.; Lewis, J. E.; Waddell, A.; Wu, S.; Steger, J. M.; Lydon, M. L.; Chait, A.; Zhao, L. Y.; Ding, H.; Li, J.-L.; Purayil, H. T.; Huo, Z.; Daaka, Y.; Garrett, T. J.; Liao, D. DAXX Drives de Novo Lipogenesis and Contributes to Tumorigenesis. Nat. Commun. 2023, 14 (1), 1927. 10.1038/s41467-023-37501-0.

(20) Wang, D.; Mahmud, I.; Thakur, V. S.; Tan, S. K.; Isom, D. G.; Lombard, D. B.; Gonzalgo, M. L.; Kryvenko, O. N.; Lorenzi, P. L.; Tcheuyap, V. T.; Brugarolas, J.; Welford, S. M. GPR1 and CMKLR1 Control Lipid Metabolism to Support the Development of Clear Cell Renal Cell Carcinoma. Cancer Res. 2024, 84 (13), 2141–2154. 10.1158/0008-5472.CAN-23-2926.

(21) Tan, S. K.; Mahmud, I.; Fontanesi, F.; Puchowicz, M.; Neumann, C. K. A.; Griswold, A. J.; Patel, R.; Dispagna, M.; Ahmed, H. H.; Gonzalgo, M. L.; Brown, J. M.; Garrett, T. J.; Welford, S. M. Obesity-Dependent Adipokine Chemerin Suppresses Fatty Acid Oxidation to Confer Ferroptosis Resistance. Cancer Discov. 2021, 11 (8), 2072–2093. 10.1158/2159-8290.CD-20-1453.

(22) Lee, B.; Mahmud, I.; Marchica, J.; Dereziński, P.; Qi, F.; Wang, F.; Joshi, P.; Valerio, F.; Rivera, I.; Patel, V.; Pavlovich, C. P.; Garrett, T. J.; Schroth, G. P.; Sun, Y.; Perera, R. J. Integrated RNA and Metabolite Profiling of Urine Liquid Biopsies for Prostate Cancer Biomarker Discovery. Sci. Rep. 2020, 10 (1), 3716. 10.1038/s41598-020-60616-z.

(23) Das, S.; Finney, A. C.; Anand, S. K.; Rohilla, S.; Liu, Y.; Pandey, N.; Ghrayeb, A.; Kumar, D.; Nunez, K.; Liu, Z.; Arias, F.; Zhao, Y.; Pearson-Gallion, B. H.; McKinney, M. P.; Richard, K. S. E.; Gomez-Vidal, J. A.; Abdullah, C. S.; Cockerham, E. D.; Eniafe, J.; Yurochko, A. D.; Magdy, T.; Pattillo, C. B.; Kevil, C. G.; Razani, B.; Bhuiyan, M. S.; Seeley, E. H.; Galliano, G. E.; Wei, B.; Tan, L.; Mahmud, I.; Surakka, I.; Garcia-Barrio, M. T.; Lorenzi, P. L.; Gottlieb, E.; Salido, E.; Zhang, J.; Orr, A. W.; Liu, W.; Diaz-Gavilan, M.; Chen, Y. E.; Dhanesha, N.; Thevenot, P. T.; Cohen, A. J.; Yurdagul, A.; Rom, O. Inhibition of Hepatic Oxalate Overproduction Ameliorates Metabolic Dysfunction-Associated Steatohepatitis. Nat. Metab. 2024, 6 (10), 1939–1962. 10.1038/s42255-024-01134-4.

(24) Lee, B.; Mahmud, I.; Pokhrel, R.; Murad, R.; Yuan, M.; Stapleton, S.; Bettegowda, C.; Jallo, G.; Eberhart, C. G.; Garrett, T.; Perera, R. J. Medulloblastoma Cerebrospinal Fluid Reveals Metabolites and Lipids Indicative of Hypoxia and Cancer-Specific RNAs. Acta Neuropathol. Commun. 2022, 10 (1), 25. 10.1186/s40478-022-01326-7.

(25) de Jong, F. A.; Beecher, C. Addressing the Current Bottlenecks of Metabolomics: Isotopic Ratio Outlier Analysis™, an Isotopic-Labeling Technique for Accurate Biochemical Profiling. Bioanalysis 2012, 4 (18), 2303–2314. 10.4155/bio.12.202.

(26) Hiller, K.; Metallo, C. M.; Kelleher, J. K.; Stephanopoulos, G. Nontargeted Elucidation of Metabolic Pathways Using Stable-Isotope Tracers and Mass Spectrometry. Anal. Chem. 2010, 82 (15), 6621–6628. 10.1021/ac1011574.

(27) Hoffmann, F.; Jaeger, C.; Bhattacharya, A.; Schmitt, C. A.; Lisec, J. Nontargeted Identification of Tracer Incorporation in High-Resolution Mass Spectrometry. Anal. Chem. 2018, 90 (12), 7253–7260. 10.1021/acs.analchem.8b00356.

(28) Mahmud, I.; Wei, B.; Veillon, L.; Tan, L.; Martinez, S.; Tran, B.; Raskind, A.; de Jong, F.; Akbani, R.; Weinstein, J. N.; Beecher, C.; Lorenzi, P. L. An IROA Workflow for Correction and Normalization of Ion Suppression in Mass Spectrometry-Based Metabolomic Profiling Data. Res. Sq. 2024, rs.3.rs-3914827. 10.21203/rs.3.rs-3914827/v1.

(29) Pinto, F. G.; Mahmud, I.; Harmon, T. A.; Rubio, V. Y.; Garrett, T. J. Rapid Prostate Cancer Noninvasive Biomarker Screening Using Segmented Flow Mass Spectrometry-Based Untargeted Metabolomics. J. Proteome Res. 2020, 19 (5), 2080–2091. 10.1021/acs.jproteome.0c00006.

(30) He, L.; Diedrich, J.; Chu, Y.-Y.; Yates, J. R. Extracting Accurate Precursor Information for Tandem Mass Spectra by RawConverter. Anal. Chem. 2015, 87 (22), 11361–11367. 10.1021/acs.analchem.5b02721.

(31) Pluskal, T.; Castillo, S.; Villar-Briones, A.; Orešič, M. MZmine 2: Modular Framework for Processing, Visualizing, and Analyzing Mass Spectrometry-Based Molecular Profile Data. BMC Bioinformatics 2010, 11 (1). 10.1186/1471-2105-11-395.

(32) Chong, J.; Xia, J. MetaboAnalystR: An R Package for Flexible and Reproducible Analysis of Metabolomics Data. Bioinformatics 2018, 34 (24), 4313–4314. 10.1093/bioinformatics/bty528.

(33) Pang, Z.; Chong, J.; Zhou, G.; de Lima Morais, D.A.; Chang, L.; Barrette, M.; Gauthier, C.; Jacques, P.-É.; Li, S.; Xia, J. MetaboAnalyst 5.0: Narrowing the Gap between Raw Spectra and Functional Insights. Nucleic Acids Res. 2021, No. gkab382. 10.1093/nar/gkab382.

(34) Xiao, J. F.; Zhou, B.; Ressom, H. W. Metabolite Identification and Quantitation in LC-MS/MS-Based Metabolomics. Trends Anal. Chem. TRAC 2012, 32, 1–14. 10.1016/j.trac.2011.08.009.

(35) Hotelling, H. Analysis of a Complex of Statistical Variables into Principal Components. J. Educ. Psychol. 19340101, 24 (6), 417. 10.1037/h0071325.

(36) Mahmud, I.; Kousik, C.; Hassell, R.; Chowdhury, K.; Boroujerdi, A. F. NMR Spectroscopy Identifies Metabolites Translocated from Powdery Mildew Resistant Rootstocks to Susceptible Watermelon Scions. J. Agric. Food Chem. 2015, 63 (36), 8083–8091. 10.1021/acs.jafc.5b02108.

(37) Gromski, P. S.; Muhamadali, H.; Ellis, D. I.; Xu, Y.; Correa, E.; Turner, M. L.; Goodacre, R. A Tutorial Review: Metabolomics and Partial Least Squares-Discriminant Analysis – a Marriage of Convenience or a Shotgun Wedding. Anal. Chim. Acta 2015, 879, 10–23. 10.1016/j.aca.2015.02.012.

(38) Haystack, a web-based tool for metabolomics research. https://www.ncbi.nlm.nih.gov/pmc/articles/PMC4251040/ (accessed 2019-10-28).

(39) Mahmud, I.; Pinto, F. G.; Rubio, V. Y.; Lee, B.; Pavlovich, C. P.; Perera, R. J.; Garrett, T. J. Rapid Diagnosis of Prostate Cancer Disease Progression Using Paper Spray Ionization Mass Spectrometry. Anal. Chem. 2021, 93 (22), 7774–7780. 10.1021/acs.analchem.1c00943.

